# Atypical tetracyclines promote longevity and ferroptotic neuroprotection via translation attenuation

**DOI:** 10.64898/2026.01.09.698733

**Authors:** Khalyd J. Clay, Manuel Sanchez-Alavez, Ian Newman, Na Na, Ana P. Verduzco Espinoza, Alan To, Shannon Saad, Hollis T. Cline, Michael Petrascheck

## Abstract

Preclinical and clinical studies have reported neuroprotective and geroprotective effects of tetracyclines that are independent of their antibiotic activity, but the underlying mechanisms remain unclear. Here, we systematically profile widely used tetracyclines, including impurities and degradation products, and identify translation attenuation as the shared driver of their neuroprotective and longevity-promoting effects, independent of classical tetracycline mechanisms. Instead, we uncover two mechanistically distinct classes of tetracyclines. Mitochondrial-targeting tetracyclines (MitoTets), exemplified by doxycycline, inhibit the mitochondrial ribosome and attenuate cytosolic translation through activation of the Integrated Stress Response (ISR). In contrast, atypical tetracyclines such as 4-epiminocycline and 12-aminominocycline act as cytosolic-targeting tetracyclines (CytoTets), directly inhibiting the cytosolic ribosome, bypassing the ISR, and protecting neurons from ferroptotic cell death. CytoTets are non-antibiotic, brain-penetrant, and neuroprotective in mouse and human neurons, establishing the tetracyclines as a tunable chemical scaffold for selectively targeting translation in aging and neurodegeneration.

**Highlights:**

- The tetracyclines broadly attenuate translation in multiple eukaryotic models
- Translation attenuation results from both ISR-dependent and ISR-independent mechanisms
- Discovery of atypical, cytosolic targeting tetracyclines (CytoTETs) that protect from ferroptosis ISR-independently
- CytoTETs inhibit translation and are neuroprotective in human-derived neurons and mouse hippocampus

## Introduction

There have long been tantalizing observations in the literature of tetracyclines eliciting anti-inflammatory and neuroprotective effects in age-associated diseases such as rheumatoid arthritis or neurodegeneration^1–9^. Similarly, tetracyclines have been found to extend lifespan in model organisms such as *C. elegans* or *D. Melanogaster*, suggesting that they have genuine geroprotective effects^10–13^. Despite clinical evidence that tetracyclines have non-antibiotic effects that are beneficial to humans, their antibiotic activity limits dosing and confounds trial results due to gastrointestinal side effects^9,14^. Consequently, non-antibiotic tetracyclines that retain their therapeutic and geroprotective profile would be desirable. However, the multitude of different tetracycline analogs, disease indications, and proposed mechanisms obscures whether tetracyclines cohesively act through one central or several distinct mechanisms, hindering the development of geroprotective compounds.

The primary antibiotic mechanism of tetracyclines is the inhibition of the bacterial ribosome, preventing translation^15^. Older studies, comparing the activity of tetracyclines in *E. coli* and *S. cerevisiae*, showed that tetracyclines target mitochondrial ribosomes in *S. cerevisiae*^16^. These findings, in light of the ancestral origin of mitochondria^17^, provide a convincing narrative for their mechanism in eukaryotes. Indeed, doxycycline was shown to target the mitochondrial ribosome in several eukaryotes^18–22^ through a mechanism that involves the activation of the mitochondrial unfolded protein response (UPR^mt^) and the Integrated Stress Response (ISR)^23–25^. However, unbiased chemoproteomics and biochemical methods have also shown that some tetracyclines directly bind to the cytosolic ribosome^11,26,27^. Inhibition of the ribosome and reducing translation is well established genetically to increase lifespan^28–32^. The most ubiquitous mechanism cited for the beneficial effects of tetracyclines observed in mammals is the inhibition of Matrix Metalloproteinase 9 (MMP9)^33–35^ originating from observations in diabetes–induced gingivitis^2^, arthritis^27^, and cancer^28^.

The non-antibiotic beneficial effects of the tetracyclines have been investigated in a variety of separate models using varying analogs, which raises the possibility that the various beneficial effects reported for different tetracyclines are the result of different mechanisms dependent on the tetracycline or the result of specific disease contexts. To elucidate the mechanism(s) by which the tetracyclines elicit their neuroprotective and geroprotective effects, we systematically profiled 21 widely used tetracyclines, including their known impurities. We established their neuroprotective and geroprotective properties using a combination of *C. elegans*, mammalian cell lines, primary neurons, and neurons derived from human induced pluripotent stem cells (iPSCs) and evaluated the validity of several proposed mechanisms of action. We demonstrate that the neuroprotective and geroprotective effects do not require inhibition of matrix metalloproteinase 9 (MMP9). Instead, we find that tetracyclines attenuate translation in eukaryotes, through both ISR-dependent and ISR-independent mechanisms, revealing an unexpected mechanistic diversity responsible for their neuroprotective and lifespan extending effects despite their close structural similarities.

## Results

### Antibiotic activity is unrelated to tetracycline-induced neuro- or geroprotection

To survey the neuroprotective and geroprotective^36,37^ properties of tetracyclines, we generated a library of the most widely studied tetracyclines, including several synthesis impurities and degradation products. To compare their neuroprotective efficacy, we measured their ability to protect hippocampal-derived HT22 cells from oxytosis/ferroptosis-mediated cell death. Oxytosis/ferroptosis is a lipid peroxidation-mediated cell death linked to neurodegenerative and other age-associated diseases. Experimentally, it is induced by high concentrations of glutamate, erastin, or RSL3^38,39^. Fourteen of the 21 tetracyclines tested were neuroprotective, exhibiting EC_50_ values of 1.5—11 µM **(Figure 1A)**.

**Figure 1:**
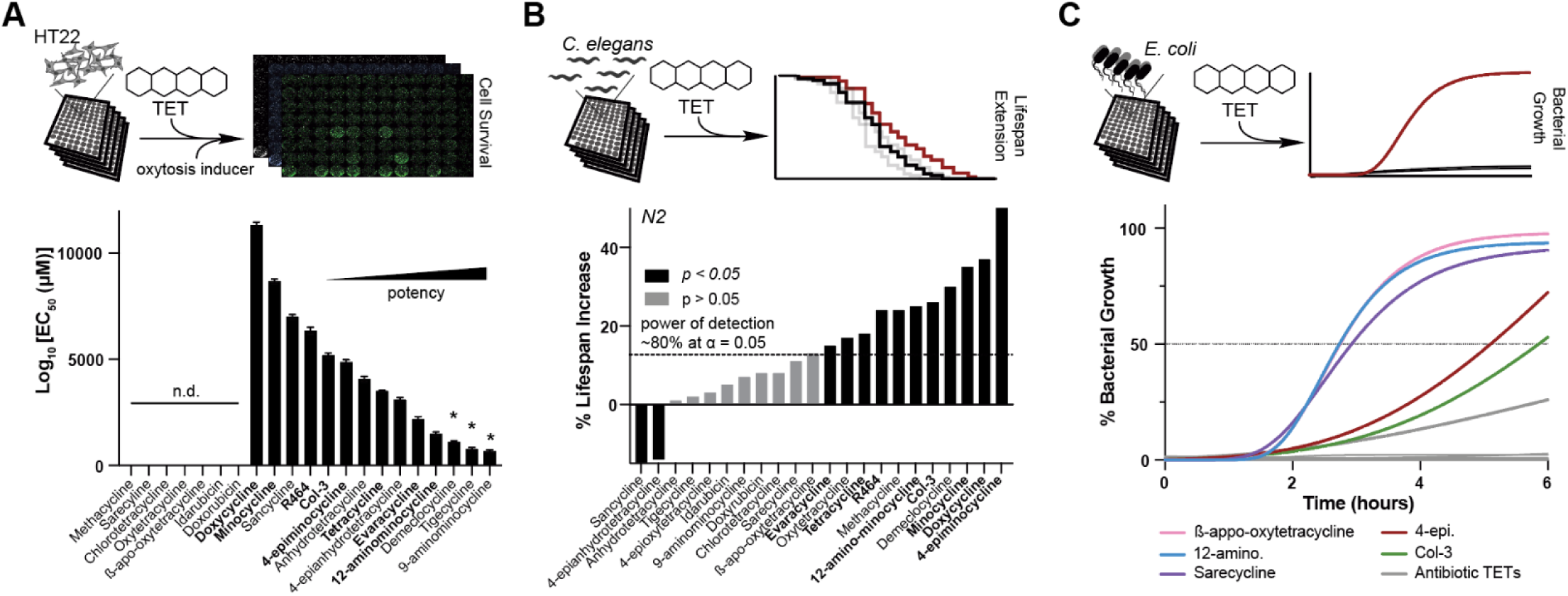
Antibiotic activity is not required for tetracycline-induced neuro- or geroprotection. Top panels depict experimental strategy to phenotypically profile 21 commercially available tetracyclines across multiple species for neuroprotection **(A),** geroprotection **(B)**, and antibiotic activity **(C)**. **(A)** Graph shows Log_10_(EC_50_) for neuroprotection. EC_50_ values were derived from dose response curves measuring cell viability of hippocampal-derived HT22 cells subjected to oxytosis/ferroptosis. HT22 cells were pre-treated for 4 hrs with increasing concentrations of tetracyclines, followed by the induction of oxytosis/ferroptosis with glutamate (5 mM) and determination of cell survival 20 hrs later. Error bars indicated mean ± SD. Data representative of three independent trials. * indicates compounds that were cytotoxic at the highest concentration. **(B)** Bar graphs show % change in lifespan for N2 *C. elegans* treated with the indicated tetracycline starting on day 1 of adulthood. Doses tested were 33 or 100 µM, except for 12-aminominocycline, which was tested at 200 μM. Significance was determined by the long-rank test. Black bars: p < 0.05; grey bars: p > 0.05. N > 50 animals per treatment. See Supplementary Data for more details. The dotted line indicates the estimated power of detection. We expect to identify 80% of all tetracyclines that extend lifespan by 15% or more at an α = 0.05**. (C)** Bacterial growth as a function of time for *E. coli* (OP50) in the presence of 100 µM of each tetracycline. Tetracyclines that allowed more than 50% growth were considered reduced-antibiotic, while tetracyclines that had the same growth curve as DMSO were considered non-antibiotic. DMSO is omitted as it is identical to that of β-appo-oxytetracyline. DMSO and kanamycin were used as negative and positive controls, respectively. Total of 3 independent experiments.

To evaluate the geroprotective properties of tetracyclines, we determined their ability to extend the lifespan of wild-type *C. elegans* (N2). We identified 11 tetracyclines that increase lifespan, eight of which were also neuroprotective **(Figure 1B, bold names; Supplementary Table 1)**. Five additional tetracyclines showed a tendency to extend lifespan but did not reach significance. These may be true negatives or their effect size might fall below our power of detection, which we estimated to be 80% at an α = 0.05 for a 15% increase in lifespan^40^ **(Figure 1B)**.

Next, we screened our library for antibiotic activity against the Gram-negative bacterium *E. coli* (**Figure 1C**). We identified five non- or reduced-antibiotic tetracyclines. These included: 12-aminominocycline and 4-epiminocycline^41^, two common minocycline impurities, β-apo-oxytetracycline, a degradation product of oxytetracycline^42^, sarecycline, a narrow-spectrum antibiotic with low activity against *E. coli*^43^, and Col-3. The two minocycline impurities, 4-epiminocycline and 12-aminominocycline, along with Col-3, also exhibited neuroprotective and geroprotective properties. Thus, the beneficial effects of tetracyclines in eukaryotes are separable from their antibiotic activity.

### MMP9 inhibition is not required for tetracycline-induced neuroprotection

Tetracycline neuroprotection is often attributed to MMP9 inhibiton via Zn^2+^ chelation^4,44–46^, but two analogs we identified (12-aminominocycline and R464) lack the C11-C12 β-diketone structure required for chelation **(Extended Figure 1A)** ^47^. We therefore tested if MMP9 inibiton was responsible for neuroprotection. As expected, they failed to chelate Zn^2+^, and even classical tetracyclines did so only weakly, with millimolar IC50 values **(Extended Figure 1B)**. In a recombinat MMP9 assays, 5 of 8 tetracyclines inhibited the enzyme at 100 µM **(Extended Figure 1C),** and dose response curves showed that 12-aminominocycline and 4-epiminocycline did not inhibit MMP9 despite being neuroprotective **(Extended Figure 1D)**. Thus, tetracycline–induced neuroprotection can be uncoupled from MMP9 inhibiton.

### Tetracyclines attenuate eukaryotic translation

We previously showed that the geroprotective effect of minocycline is the result of attenuated translation^11^. Excluding MMP9 inhibition as the primary mechanism raised the question of whether the attenuation of translation drives neuroprotection as well. We therefore screened our tetracycline library for the reduction of *de novo* translation in HEK293 cells through o-propargyl puromycin incorporation (OPP)^48^. The OPP-labeled proteins are visualized by conjugating a fluorophore using click chemistry^49^, and quantified by comparing the green probe signal to the nuclear DAPI signal. Of the 21 tetracyclines, 17 reduced *de novo* translation in HEK293 cells (**Figure 2A**). The efficacy of tetracyclines in reducing translation is weaker than that of cycloheximide. Thus, we refer to the reduction in translation by tetracyclines in eukaryotes as *attenuation*. Our data show attenuation of translation is a general feature of tetracyclines in eukaryotes, contradicting the long-held belief that tetracyclines specifically inhibit translation in bacteria.

**Figure 2:**
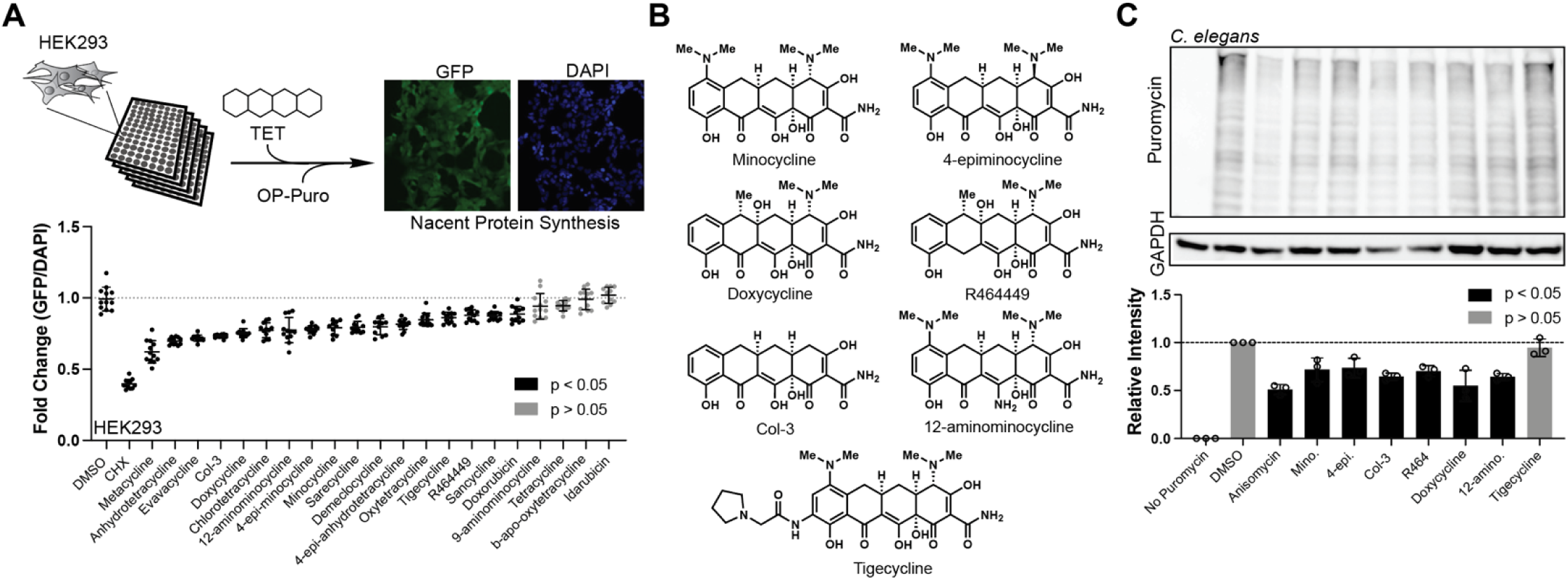
Tetracyclines attenuate translation in eukaryotes. **(A)** Graph shows fold change in *de novo* translation in HEK293 cells treated with individual tetracyclines. Each tetracycline was tested at a concentration of 100 µM. Translation was quantified by the OPP incorporation assay determining the mean OPP incorporation-intensity relative to DAPI. Cycloheximide (CHX) serves as a positive control. Total of 3 independent trials with at least 4 images quantified per trial. **(B)** Chemical structures of 7 diverse tetracyclines, representing an array of structural modifications. **(C)** A puromycin incorporation assay (SUnSET) was used to monitor the effect of tetracycline treatment on translation in *C. elegans* (100 µM, for 4 hrs). Immunoblot of protein extracts stained with puromycin antibody reveals reduced puromycin incorporation and hence translation (top). GAPDH is used as a loading control. Quantification of three independent SUnSET trials shows broad translation inhibition of most tetracyclines. Significance was determined by one-way ANOVA using Dunnett’s multiple comparisons correction, where black represents p < 0.05 and grey represents p > 0.05.

We further determined if all geroprotective tetracyclines attenuate translation in wild-type *C. elegans*. We selected 6 geroprotective tetracyclines and one inactive negative control (tigecycline) **(Figure 2B)** and measured translation in tetracycline-treated *C. elegans* using the SUnSET protocol, which visualizes the incorporation of puromycin into newly synthesized proteins via immunoblot^50,51^. All 6 geroprotective tetracyclines attenuated translation, while tigecycline did not **(Figure 2C)**. Notably, tigecycline attenuated translation and exhibited neuroprotective effects in mammalian cells, suggesting that its binding site is sufficiently divergent between species. Thus, the attenuation of translation by tetracyclines is likely to be responsible for their geroprotective effects.

### Tetracyclines elicit neuroprotection by ISR-dependent and independent attenuation of translation

We next set out to characterize the mechanisms by which tetracyclines attenuate translation. Tetracyclines, like doxycycline, that target the mitochondrial ribosome activate the ISR via the phosphorylation of eIF2*α* to attenuate cytosolic translation, and activate ATF4^18,20–22^. Alternatively, tetracyclines that target the cytosolic ribosome directly attenuate translation, independently of the ISR^11^. The two mechanisms can be distinguished by co-treatment with the ISR Inhibitor (ISRIB)^52,53^, which prevents tetracycline-mediated attenuation of translation in cells treated with ISR-dependent tetracyclines, but not in cells treated with ISR-independent tetracyclines.

To determine if the tetracycline-mediated attenuation of translation is prevented by co-treatment with ISRIB, we repeated the OPP-based translation assay in HT22 cells in the presence or absence of ISRIB. As a positive control for an ISR-*dependent* inhibition of translation, we included thapsigargin^54^. As a positive control for the ISR-*independent* inhibition of translation, we included cycloheximide (CHX)^55^.

These experiments revealed the existence of two tetracycline classes, ISR-dependent (**Figure 3A, C**) and ISR-independent tetracyclines(**Figure 3B, C**). The ISR-dependent tetracyclines, such as doxycycline, were previously shown to act on the mitochondrial ribosome^18,22^ while the ISR-independent tetracyclines, such as minocycline, where shown to directly bind to the cytoplasmic ribosome to lower translation^11,23^. We thus will refer to mitochondrial targeting tetracyclines as MitoTets (e.g. doxycycline) and to those targeting the cytoplasmic ribosome (minocycline derivatives) as CytoTets. The ability of MitoTets such as doxycycline to attenuate translation can be blocked by ISRIB, allowing us to directly test whether the neuroprotection depends on the attenuation of translation. Co-treatment of doxycycline-treated cells with ISRIB rescued translation and abolished the neuroprotective effect, indicating that attenuation of translation leads to neuroprotection.

**Figure 3:**
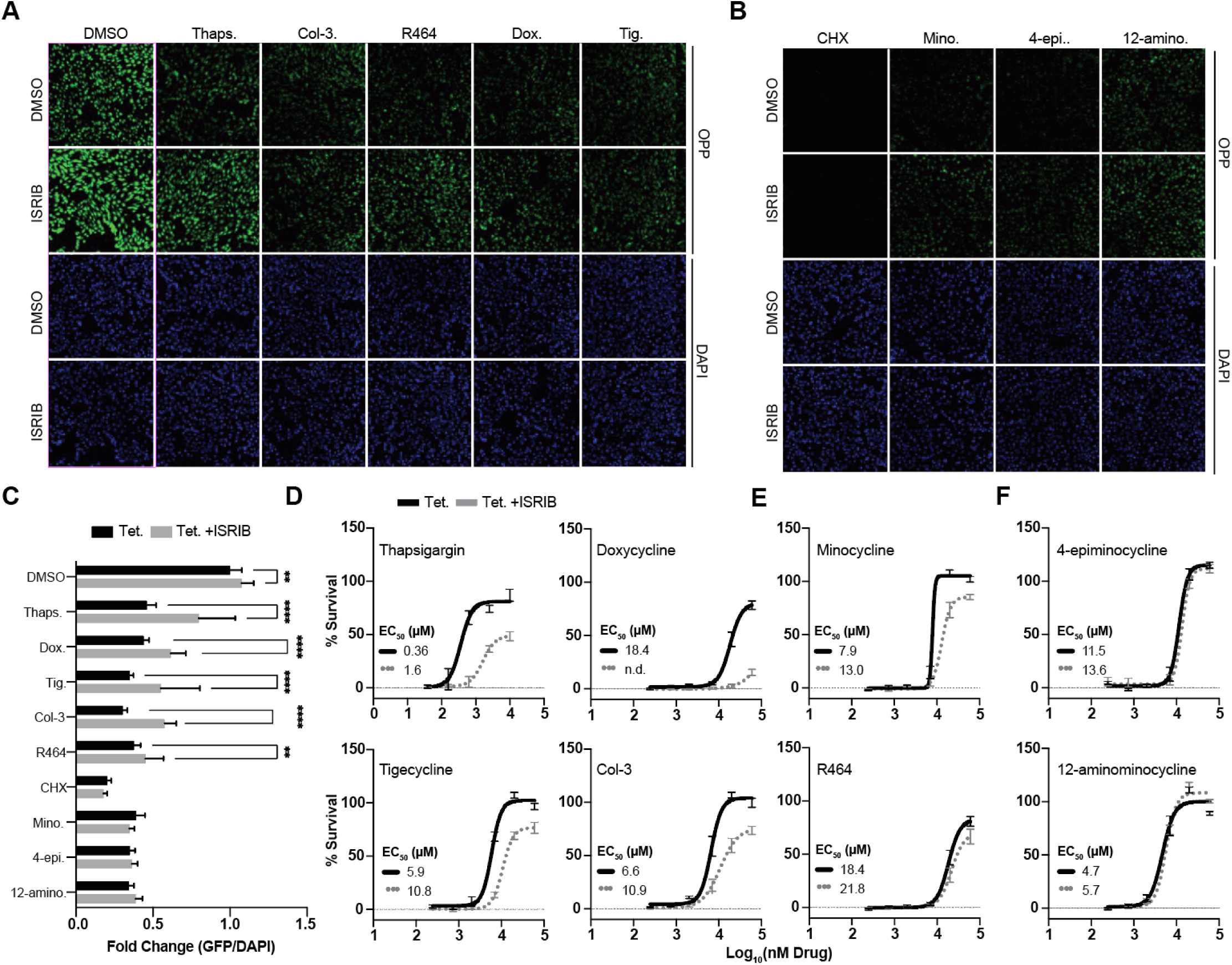
Tetracyclines protect neurons by ISR-dependent and ISR-independent attenuation of translation. **(A)** Representative fluorescent micrographs of HT22 cells stained for OPP incorporation as a readout for *de novo* translation. Translation is attenuated by ISR-dependent tetracyclines (MitoTets) and restored by co-treatment with ISRIB. Thapsigargin (Thaps., 1 μM) is included as a control for an ISR-dependent translation inhibitor. Tetracyclines were screened at 100 μM. **(B)** Same fluorescent micrographs as in **A,** but for ISR-independent tetracyclines (CytoTets) whose translation attenuation is unaffected by ISRIB co-treatment. Cycloheximide (CHX, 500 nM) is included as a control for an ISR-independent translation inhibitor. **(C)** Quantification of the OPP incorporation-intensity relative to the DAPI signals shown in Figure 4A & 4B. Several tetracyclines attenuate translation independently of the ISR (black vs. grey bars). Significance was determined by two-way ANOVA with Šídák multiple comparisons test. ** = p < 0.01 and **** = p < 0.0001. **(D)** Graphs show the % survival of HT22 cells as a function of tetracycline dose. The % survival is calculated relative to non-glutamate-treated HT22 control cells (100%). HT22 cells were pre-treated with ISR-dependent tetracyclines (MitoTets), alone or in combination with ISRIB followed by the induction of oxytosis/ferroptosis. Thapsigargin (Thaps.) is used as a positive, ISR-dependent control. **(E)** Same as **D,** but for partially ISR-dependent tetracyclines. **(F)** Same as **D,** but for ISR-independent (CytoTets) tetracyclines. For all figures: Cells were treated for 2 hrs with ISRIB (300 nM, grey bar, or dotted line) or vehicle control (DMSO, black bar/line) prior to co-incubation with the indicated tetracycline or control compound. Error bars indicate mean ± SD from three independent trials.

Of the 7 tetracyclines, we classified doxycycline, Col-3, and tigecycline as MitoTets (**Figure 3D**); Minocycline and R464 as intermediate (**Figure 3E**); and 4-epiminocycline and 12-aminominocycline as CytoTets (**Figure 3F**). Doxycycline is the clearest MitoTet representative, as ISRIB co-treatment reduced its neuroprotective properties the most. 4-epiminocycline and 12-aminominocycline were the clearest CytoTet representatives and entirely resistant to ISRIB co-treatment (**Figure 3D, F**). However, ISRIB co-treatment never completely rescued translation or completely abolished neuroprotection, even for MitoTets, suggesting that all tetracyclines inhibit the cytosolic ribosome to varying degrees and that ISR activation occurs in addition. In summary, tetracyclines diverge into MitoTets and CytoTets based on the mechanism underlying their neuroprotective effect. We were unable to pinpoint unequivocal structural motifs responsible for the divergence between MitoTets and CytoTets, but modification in the central pharmacophore important to antibiotic activity (e.g., C11-12) tended to result in ISR-independent CytoTets. Similarly, inversion of the stereochemistry at the C4 position reduced antibiotic activity and increased potency.

### Tetracyclines extend lifespan by ISR-dependent and independent mechanisms in *C. elegans*

We next asked if the classification of tetracyclines into MitoTets and CytoTets also applies to their geroprotective effects. Consistent with this classification, the MitoTet doxycycline is known to inhibit the mitochondrial ribosome, triggering the mitochondrial unfolded protein response (UPR^mt^), which in turn activates the ISR to attenuate translation and upregulate ATF-4 to extend the lifespan of *C. elegans*^56–58^. Conversely, CytoTets are expected to extend lifespan independently of the ISR or of any stress response, such as the heat shock response (HSR) or the UPR^mt^, as stress response signaling converges on the ISR^59^. Whether tetracyclines exist that extend lifespan in *C. elegans* independently of the ISR, as seen for the CytoTets in cell culture, is unknown.

We previously showed that hyper-translation is a defining phenotype of the HSR-deficient *hsf-1(sy441)* mutant^11,60^. Hyper-translation makes the *hsf-1(sy441)* mutant especially resistant to many longevity paradigms, including UPR^ER^ activation, reduced of mTOR activity, reduced IIS signaling, dietary restriction or hormesis^61–65^, while sensitizing animals to interventions that lower translation. Thus, we tested 21 tetracyclines for their ability to extend the lifespan of *hsf-1(sy441)* mutants^11,60^. Eleven tetracyclines extended the lifespan of *hsf-1(sy441)* (**Figure 4A**). These 11 tetracyclines included both MitoTets and CytoTets, based on their classification in mammalian cells. Thus, both MitoTets and CytoTets extend lifespan independently of the HSR by lowering the excessive protein synthesis of the *hsf-1(sy441)* mutant. This conclusion was further corroborated by evaluating the induction of the *hsp-16.2::GFP* heat shock reporter after treatment with 2 MitoTets and 2 CytoTets, followed by a heat shock. Both MitoTets and CytoTets inhibit rather than activate the HSR **(Extended Figure 4A—4C).**

**Figure 4:**
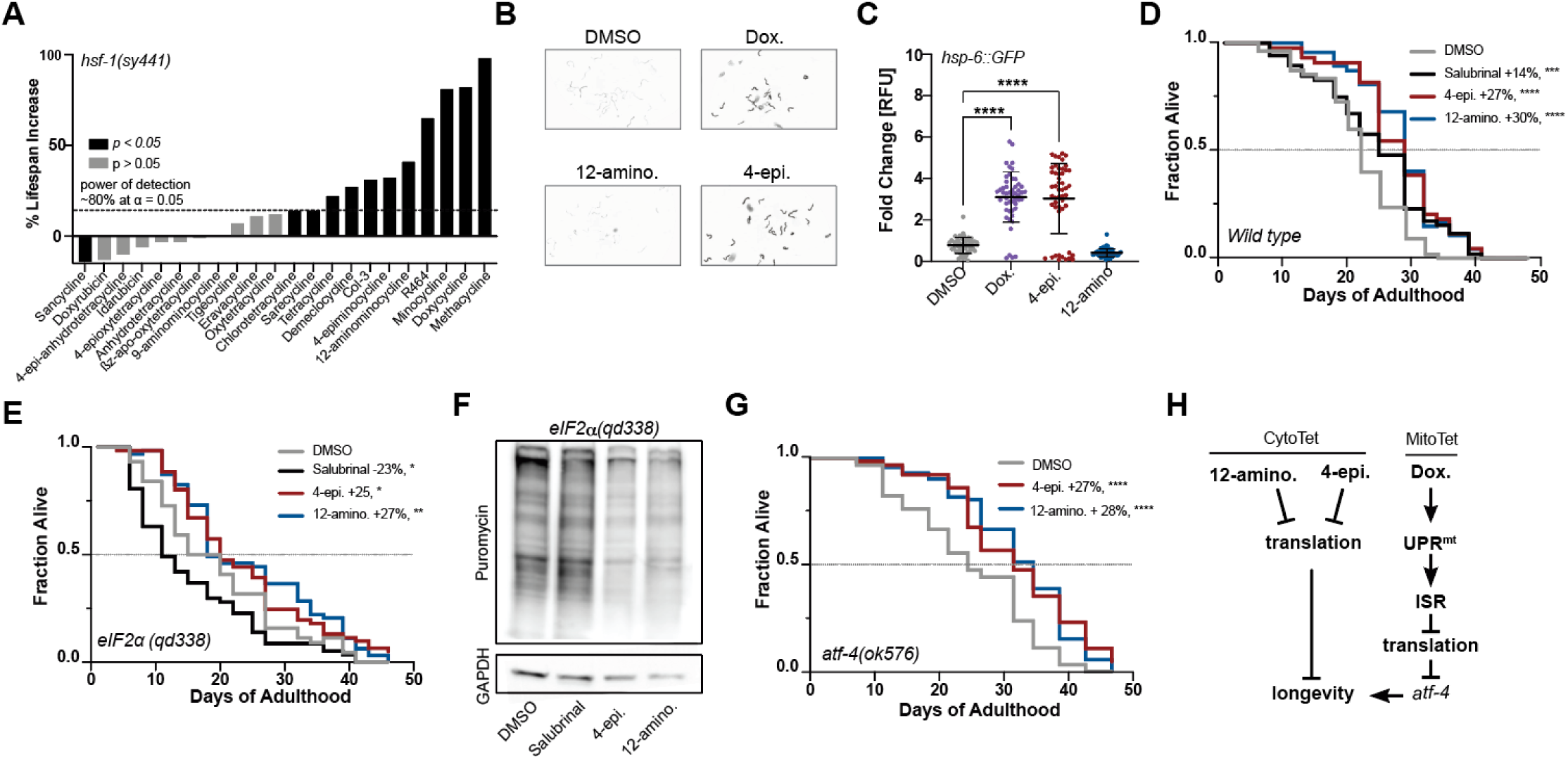
4-epiminocycline and 12-aminocycline extend lifespan independently of the ISR. **(A)** Bar graphs show % change in lifespan for *hsf-1(sy441) mutants* treated with the indicated tetracycline starting on day 1 of adulthood. Doses tested were 33 or 100 µM, except for 12-aminominocycline, which was tested at 200 μM. Black bars p < 0.05, grey bars p > 0.05. N > 50 animals per treatment. See Supplementary Data for more details. The dotted line indicates the estimated power of detection. We expect to identify 80% of all tetracyclines that extend lifespan by 15% or more at an α = 0.05. For additional data on the HSR, see Extended Figure 5A—5C. **(B)** Representative fluorescent images of 50 randomly selected *hsp-6p::GFP* UPR^mt^ reporter animals treated with the indicated tetracycline. Images were taken in parallel following measurement in Figure 5C. Images were inverted for clarity. **(C)** Scatter plot shows the fold induction relative to DMSO of the *hsp-6::GFP* UPR^mt^ reporter in response to 12-hrs tetracycline treatment initiated at the L2 stage. Significance was determined by one-way ANOVA with Dunnett’s multiple comparisons, where **** = p < 0.0001. Error bars indicate mean ± SD of 3 independent trials. **(D)** Survival plot of wild-type (N2) animals treated with tetracyclines or the ISR-dependent translation inhibitor salubrinal. Tetracycline and salubrinal treatment extend the lifespan of N2. **(E)** Survival plot of ISR-deficient *eIF2⍺(qd338)* mutants treated with tetracyclines or salubrinal. Only tetracyclines, but not salubrinal treatment, extend the lifespan of *eIF2⍺(qd338)* mutants. For similar data on *eIF2⍺(rog3),* see Extended Figure 5D. **(F)** A puromycin incorporation assay (SUnSET) was used to monitor the effect of tetracycline or salubrinal treatment on translation in the ISR-deficient *eIF2⍺(qd338)* mutant. Only tetracyclines, but not salubrinal treatment, reduce translation in the ISR-deficient *eIF2⍺(qd338)* mutant. GAPDH was used as a loading control. For quantification, see Extended Figure 5E. **(G)** Survival plot of the partially ISR-deficient *atf-4(ok576)* mutants treated with tetracyclines. Tetracyclines extend lifespan mostly independent of ATF-4. For corresponding survival data on doxycycline, see Extended Figure 5F & 5G. **(H)** Schematic outlining distinguishing features between CytoTets and MitoTets. CytoTets, as exemplified by 12-aminominocycline and 4-epiminocycline, directly inhibit translation to extend lifespan, while MitoTets, as exemplified by doxycycline, activate the mitochondrial UPR to inhibit translation and subsequently extend lifespan in an *atf-4* dependent manner. The significance of all survival data was determined by the log-rank test. In **B–G** all compounds were tested at 100 µM or 200 µM for 12-aminominocycline.

To probe the involvement of the UPR^mt^, we evaluated the ability of the most typical MitoTet, doxycycline, and the most typical CytoTets, 4-epiminocycline and 12-aminominocycline, to induce the UPR^mt^ reporter *hsp-6::GFP*^66^. Consistent with their classification as MitoTets and CytoTets, doxycycline induced the UPR^mt^ while 12-aminominocycline did not. Surprisingly, 4-epiminocycline induced the UPR^mt^ (**Figure 4B & C**).

The induction of the UPR^mt^ by 4-epiminocycline suggested it might act as a MitoTet in *C. elegans* but as a CytoTet in mammals. To classify 4-epiminocycline we determined its dependence on the ISR to extend lifespan. We measured the ability of 4-epiminocycline and 12-aminominocycline, to extend the lifespan of the ISR-deficient *eIF2⍺(rog3)*^67^ and *eIF2⍺(qd338)*^68^ strains. These strains carry phospho-deficient S49A and S49P mutations that prevent *eIF2⍺* phosphorylation and thus ISR activation. As a control, we included salubrinal, an ISR-dependent translation initiation inhibitor that extends the lifespan of *C. elegans*^60,69^. 4-epiminocycline and 12-aminominocycline treatment extended the lifespan of ISR-deficient strains (**Figure 4D, E, Extended Figure 4D**) and attenuated translation in the *eIF2α(qd338)* mutant **(Figure 4F, Extended Figure 4E)**, while salubrinal treatment did neither. We conclude that 4-epiminocycline and 12-aminominocycline extend lifespan and attenuate translation independently of the ISR.

To further corroborate 4-epiminocycline and 12-aminominocycline as CytoTets and of doxycycline as a MitoTet, we determined their dependency on ATF-4, an ISR transcription factor downstream of *eIF2⍺* phosphorylation. Both 4-epiminocycline and 12-aminominocycline extended the lifespan of *atf-4(ok576)* mutants **(Figure 4G),** while the effect of doxycycline was severely blunted **(Extended Figure 4F & G)**. Thus, the mechanistic classification of 4-epiminocycline and 12-aminominocycline as CytoTets and of doxycycline as a MitoTet applies to geroprotection as well (**Figure 4H**).

### The CytoTet 4-epiminocycline is neuroprotective and inhibits translation in mouse and human neurons

Previous studies have shown that inhibiting the ISR can improve age-related cognitive decline in mice^70^ and extend lifespan in *C. elegans*^71^. ISR-independent, non-antibiotic neuroprotective tetracyclines would make attractive therapeutics for treating aging and age-related diseases in cases where continued ISR activation may be problematic.

Although the CytoTet profile of 12-aminominocycline was cleaner than that of 4-epiminocycline, as it had no MMP9 activity and did not activate the UPR^mt^ in *C. elegans*, we focused on 4-epiminocycline as 12-aminominocycline was less stable. The pharmacokinetic (PK) evaluation of 4-epiminocycline in rats established significant exposure in both the periphery and CNS. A single 50 mg/kg dose (i.p) resulted in plasma and brain concentrations of 1887 ng/mL and 410 ng/g, respectively, while a 25 mg/kg dose resulted in plasma and brain concentrations of 764 ng/mL and 172 ng/g, respectively **(Figure 5A),** 8 hours post-injection **(Figure 5B)**. Thus, 4-epiminocycline is brain-penetrant and has a low clearance in both plasma and the CNS.

**Figure 5:**
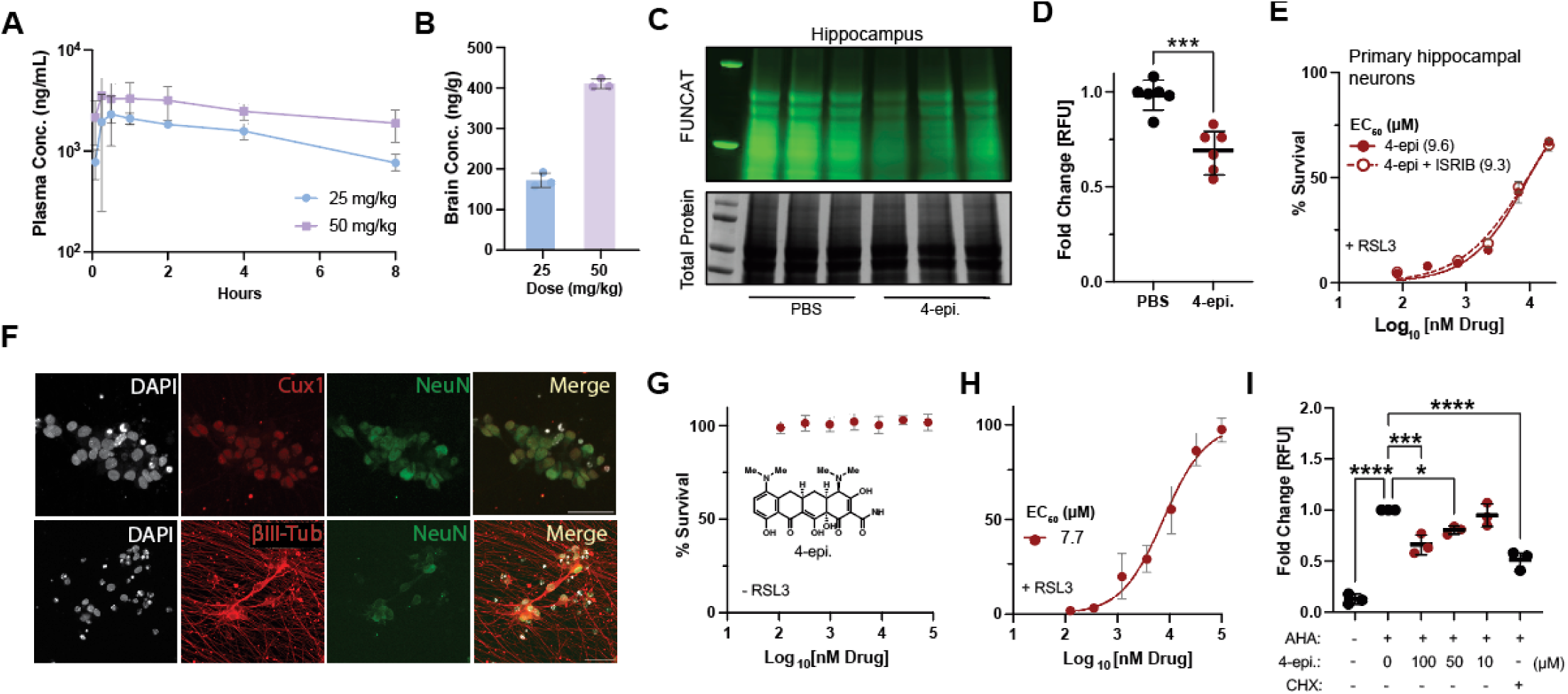
4-epiminocycline is a non-antibiotic brain penetrant CytoTet that protects human neurons from ferroptosis. **(A)** Plasma concentrations of 4-epiminocycline over 8 hrs following a single intraperitoneal injection (i.p.) of male Sprague–Dawley rats. **(B)** Brain concentrations of 4-epiminocycline after 8 hrs following a single intraperitoneal injection of male Sprague–Dawley rats. **(C)** Fluorescent-scan (top) and Coomassie blue stain (bottom) gel. In-gel fluorescence detects *de novo* translation by the amount of incorporated fluorescently labelled AHA (top) relative to total protein (bottom). 4-epiminocycline reduced *de novo* translation in the hippocampus of mice. **(D)** Quantification of FUNCAT experiments shows 4-epiminocycline reduces *de novo* translation by 25%. Significance was determined by a two-tailed Student’s t-test, where *** = p < 0.001 and N = 6. **(E)** Survival of primary hippocampal neurons as a function of 4-epiminocycline concentration after ferroptosis induction by RSL3 (300 nM). Co-treatment with ISRIB (300 nM) does not abolish the neuroprotective effect of 4-epiminocycline, confirming its classification as a CytoTet. **(F)** (Top) Cropped representative confocal images of immunofluorescent labeling of generated iNs. Nuclei (DAPI, white), cortical marker CUX1 (Cux1, red), and neuronal marker (NeuN, green) stains are shown. (Bottom) Validation of differentiation into iNeurons, by staining for nuclei (DAPI, white), the pan neuronal marker TUBB3 (βIII-Tub, red), and the neuronal marker (NeuN, green). **(G)** No cytotoxicity was detected in iNeurons 48 hrs after 4-epiminocycline addition. **(H)** Survival of iNeurons as a function of 4-epiminocycline concentration after ferroptosis induction by RSL3 (2 μM). **(I)** Quantification of *de novo* translation in iNeurons treated with increasing concentration of 4-epiminocycline using FUNCAT. iNeurons were co-incubated with (AHA, 8mM) and increasing concentrations of 4-epiminocycline for 4 hrs, followed by lysis and bioconjugation of an Alexa-488 alkyne to the incorporated AHA by a click reaction. The protein synthesis inhibitor cycloheximide (CHX, 250nM) was used as a positive control. Significance was determined by one-way ANOVA with Dunnett’s multiple comparisons, where * = p < 0.05, *** = p < 0.001 and **** = p < 0.0001. All error bars indicate mean ± SD from 3-6 independent trials or animals. Cell survival was measured using Cell Titer Glo 20 hrs after treatment in all experiments.

We then measured the ability of 4-epiminocycline to attenuate translation in a mouse brain *in vivo* by Fluorescent Non-canonical Amino Acid Tagging (FUNCAT). FUNCAT measures the incorporation of azidohomoalanine (AHA) into newly synthesized proteins and visualization by bioconjugation of a TAMRA-alkyne using click chemistry **(Extended Figure 5A)**^72^. We treated male C57BL/6 mice with 4-epiminocycline in the drinking water for three days, followed by an intraperitoneal (i.p.) co-injection with 4-epiminocycline plus AHA to label newly synthesized proteins. Since neuronal activity promotes translation^72^, we injected the mice with seizure-inducing pentylenetetrazol (PTZ) 30 minutes after the AHA injection. Two hours post-PTZ injection, the mice were sacrificed, and hippocampal proteins were extracted to quantify translation. Treatment with 4-epiminocycline significantly attenuated hippocampal translation by ∼25% **(Figure 5C & D)**, indicating that 4-epiminocycline inhibits translation *in vivo* to the same degree as observed in all other paradigms.

We further determined whether 4-epiminocycline acts as a CytoTet or MitoTet *in vivo*. The MitoTet doxycycline disrupts the mitonuclear translational balance, which can be visualized by the stoichiometric ratio between nuclear-encoded electron transport chain components (ETC), such as ATP5A, and mitochondrial-encoded ETC components like MT-CO1^12^. We did not observe any selective suppression of the mitochondrial-encoded MT-CO1, as would be expected for a MitoTet, indicating that 4-epiminocycline does not inhibit mitochondrial translation **(Extended Figure 5B & C)**. It is essential to note that we normalized each lane by protein amount and the loading control actin. This method obfuscates a cytosolic ribosome-specific inhibitor since both Actin and ATP5A are equally reduced. Taken together, these data show that 4-epiminocycline is a CytoTet and not a MitoTet like doxycycline.

We then pre-treated primary mouse neurons with 4-epiminocycline and measured survival following induction of oxytosis/ferroptosis^73^ in the presence and absence of ISRIB. As expected for an ISR-independent CytoTet, the dose-dependent improvement of survival of 4-epiminocycline-treated neurons was unaffected by ISRIB co-treatment **(Figure 5E)**. ISRIB activity was independently confirmed in primary neurons by blocking tunicamycin-induced translational repression **(Extended Figure 5D)**^54,74^.

Finally, we validated the ability of 4-epiminocycline to protect human neurons from oxytosis/ferroptosis. We used human (iPSC) line HE463#7 to differentiate into neurons (iNeurons) **(Extended Figure 5E)**^75^. We confirmed the neuronal identity of iNeurons by expression of the neuronal markers NeuN, Cux1, and TUBB3 **(Figure 5F)**. We also confirmed that 4-epiminocycline was nontoxic to iNeurons **(Figure 5G)**. We then pre-treated iNeurons with 4-epiminocycline and induced oxytosis/ferroptosis with RSL3. 4-epiminocycline dose-dependently increased the survival of human iNeurons **(Figure 5H)**. We then measured translation using FUNCAT and found a dose-dependent attenuation of translation by a maximum of 40% at the highest concentration **(Figure 5I)**. Together, we conclude that non-antibiotic CytoTets are brain penetrant and neuroprotective in mammals, providing an interesting new strategy for therapeutic development.

## Discussion

The neuroprotective, anti-inflammatory, and geroprotective effects of tetracyclines are well-documented in the pre-clinical and clinical trial literature including for many age-associated diseases. While the literature on the potential therapeutic effects of different tetracyclines is extensive, no systematic study has been conducted to compare and evaluate the underlying molecular mechanisms. Here, we profiled the most widely used tetracyclines and their impurities to characterize their geroprotective and neuroprotective activities and mechanisms of action.

Our data establish: (i) Modifications in the C4 or C11,12 positions of tetracyclines weaken their antibiotic and MMP9 inhibitory activity and uncouple their neuroprotective and geroprotective effects from antibiotic activity (**Figure 1 & Extended Figure 1**). (ii) Instead, the neuroprotective and geroprotective effects of tetracyclines are the result of attenuation of translation (**Figures 2 & 3**). (iii) The mechanism of action by which tetracyclines attenuate translation classifies them into ISR-dependent (MitoTets) and ISR-independent tetracyclines (CytoTets) (**Figures 3 & 4**). MitoTets target the mitochondrial ribosome, resulting in ISR-dependency, while CytoTets target the cytosolic ribosome directly. (iv) Minocycline degradation products like 4-epiminocycline show reduced antibiotic activity, are brain penetrant, and protect both mouse and human neurons from oxytosis/ferroptosis (**Figure 5**).

We first uncoupled the neuroprotective effects from MMP9 inhibition by showing that structurally related minocycline analogs fail to inhibit MMP9 but remain neuroprotective (**Extended Figure 1, 3E, F**). This evidence challenges the pervasive assertion that MMP9 inhibition is the causal mechanism underlying tetracycline neuroprotection, consistent with past failures of specific MMP inhibitors to recapitulate the properties of tetracyclines.

We then focused on 7 diverse tetracyclines to evaluate the attenuation of translation as the underlying mechanism for neuroprotection^11–13,76^. All 7 neuroprotective tetracyclines lower protein synthesis but did so through ISR-dependent (MitoTets) and ISR-independent (CytoTets) mechanisms (**Figure 3**). The ISR dependency of doxycycline enabled us to demonstrate that the attenuation of translation directly causes neuroprotection, since the ISR inhibitor ISRIB restored translation and abolished neuroprotection during doxycycline treatment, revealing a causal relationship (**Figure 3**). The ISR-based classification of tetracyclines into MitoTets and CytoTets also applies to their effect on longevity. **(Figure 2)**^77^. MitoTets, such as doxycycline, induce the UPR^mt^ and ISR and require ATF-4 to extend lifespan^18–22,25,56^. Conversely, CytoTets such as 12-aminominocycline and 4-epiminocycline neither require the ISR nor ATF-4 (**Figure 4**). However, our data also suggest that many tetracyclines fall in between the MitoTet and CytoTets extremes and are partially ISR dependent.

Many antibiotics that inhibit bacterial ribosomes also inhibit mitochondrial ribosomes by identical mechanisms^78,79^. Cryo-EM structures of tigecycline bound to the mitoribosome showed binding in the peptidyl transferase center that overlapped with the binding site for structurally related tetracyclines in bacterial ribosomes^19^. A recent screen aimed at identifying mitochondrial-targeting tetracyclines with limited antibiotic activity identified 9-t-butyl-doxycycline as a therapeutically developable compound for influenza^19^. Notably, none of the 18 tetracyclines derived from the minocycline scaffold activate the UPR^mt^, confirming that doxycycline-related tetracyclines tend to target mitochondrial ribosomes, whereas minocycline-related tetracyclines tend to target cytosolic ribosomes^11^. Thus, tetracyclines represent a molecular scaffold whose neuroprotective and geroprotective properties can be tuned to be ISR-dependent or independent by either targeting the mitochondrial or cytosolic ribosome.

We examined the potential of the CytoTet 4-epiminocycline as a neuroprotective agent. 4-epiminocycline is the epimerization product of minocycline and only differs in the stereochemistry of the C4 position, resulting in low antibiotic activity, while remaining brain penetrant and improving neuroprotection efficacy (**Figures 1 & 5**). 4-epiminocycline attenuated translation in the hippocampus of mice as well as in human iNeurons by ∼25% (**Figures 5D & 5I**)^80^. Translation was reduced without disrupting the mito-nuclear balance of ETC proteins **(Extended Figure Figure 5B),** as would be expected for doxycycline but not minocycline derivatives^12^. Importantly, 4-epiminocycline protected primary mouse and human iNeurons from RSL3-induced oxytosis/ferroptosis **(Figure 5E & 5H)** independent of the ISR^18^.

Minocycline, doxycycline, and other tetracyclines have been tested for their beneficial effects across various indications related to aging but unrelated to their antibiotic activity^59^. However, the results in human clinical trials have been mixed^9^. One of the critical limitations mentioned in many studies is the antibiotic activity, which is both dose-limiting and a liability for chronic use due to associated gastrointestinal side-effects and the important role of the microbiome in aging ^9,14,81,82^ Several recent studies, including this one, have demonstrated that the beneficial pre-clinical effects can be uncoupled from the antibiotic activity, as illustrated by 4-epiminocycline. In addition, the surprising mechanistic diversity of tetracyclines has clear implications for the interpretation of past and future clinical trials. The results of both failed and successful trials need to be reconsidered in light of the ISR dependency of individual tetracyclines. Activation of the ISR can have both beneficial and detrimental effects, depending on the disease indication, and thus drive the failure or success of a trial. Similarly, when generating novel non-antibiotic tetracycline derivatives, it will be crucial to consider the ISR dependency of the parent scaffold and to match it in the non-antibiotic derivatives. Finally, our identification of minocycline impurities with superior efficacy and potency may explain at least some of the mixed or disappointing results in clinical studies, as it represents a variable that was poorly controlled for in any study. Pharmacologically active metabolites with higher potency than the parent compound are well-documented in drug discovery^83^. Overall, our study suggests that tetracyclines, with their extensive safety profile offer the possibility to attenuate translation in humans and thus chemically target one of the most potent lifespan extending mechanisms identified in model organisms.

## Acknowledgments

Some strains were provided by the CGC, funded by the NIH Office of Research Infrastructure Programs (P40 OD010440). This work was supported by grants from the NIH (1RF1AG079517-01 to H.T.C. and 1R21NS107951-01 to M.P.), a Fellowship from the Helen Dorris Foundation to K.C.

---

**Extended Figure 1:**
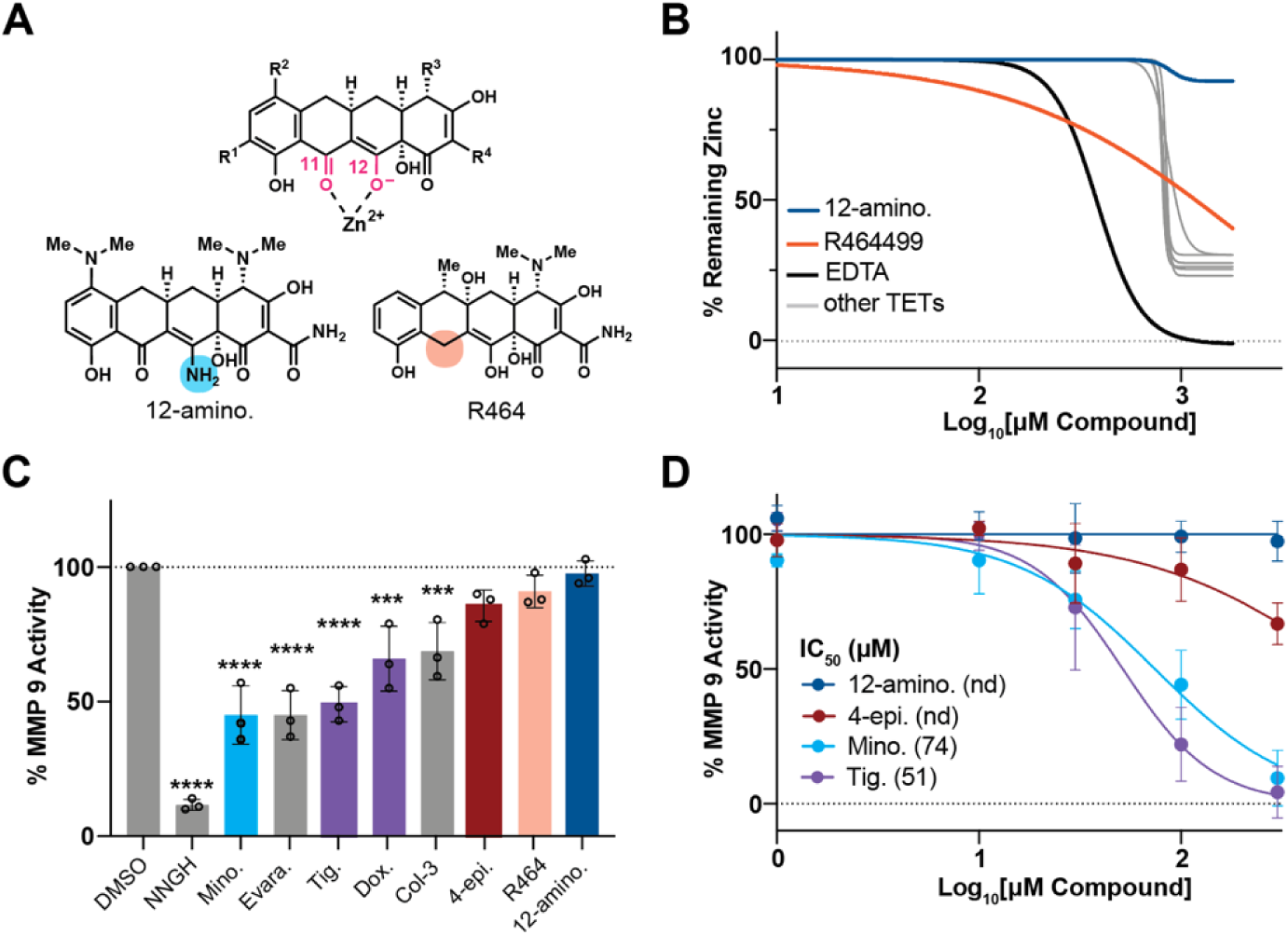
MMP 9 inhibition is not required for tetracycline-induced neuroprotection. **(A)** Shown is the general structure of the tetracyclines with the keto-enol system (pink) at C11 & C12 that is responsible for the chelation of Zn^2+^ and other divalent ions. Structural modifications in 12-aminominocycline and R464 disrupt the chelation center (blue & orange circles). **(B)** Colorimetric assay measuring the % remaining Zn^2+^ as a function of tetracycline concentration, indicative of the chelation effect of each tetracycline. 12-aminominocycline does not chelate ions at concentrations up to 1800 µM, whereas 300 µM is required for R464. Other tetracyclines tested (grey) include: 4-epiminocycline, minocycline, Col-3, Doxycycline, Tigecycline, & Evaracycline. EDTA was used as a positive control. **(C)** Bar graph shows the % remaining recombinant MMP9 activity after tetracycline treatment (100 µM). The assay measures proteolytic cleavage of a fluorogenic substrate, released upon cleavage. NNGH is a broad-spectrum inhibitor of matrix metalloproteinases and was used as a positive control. Significance was determined by one-way ANOVA with Dunnett’s multiple comparisons, where *** = p < 0.001 and **** = p < 0.0001. Error bars indicate mean ± SD from three independent trials. **(D)** Dose response curve of four neuroprotective tetracyclines. Tetracyclines with IC_50_ greater than 300 µM are indicated as “not determined” (n.d.).

**Extended Figure 4:**
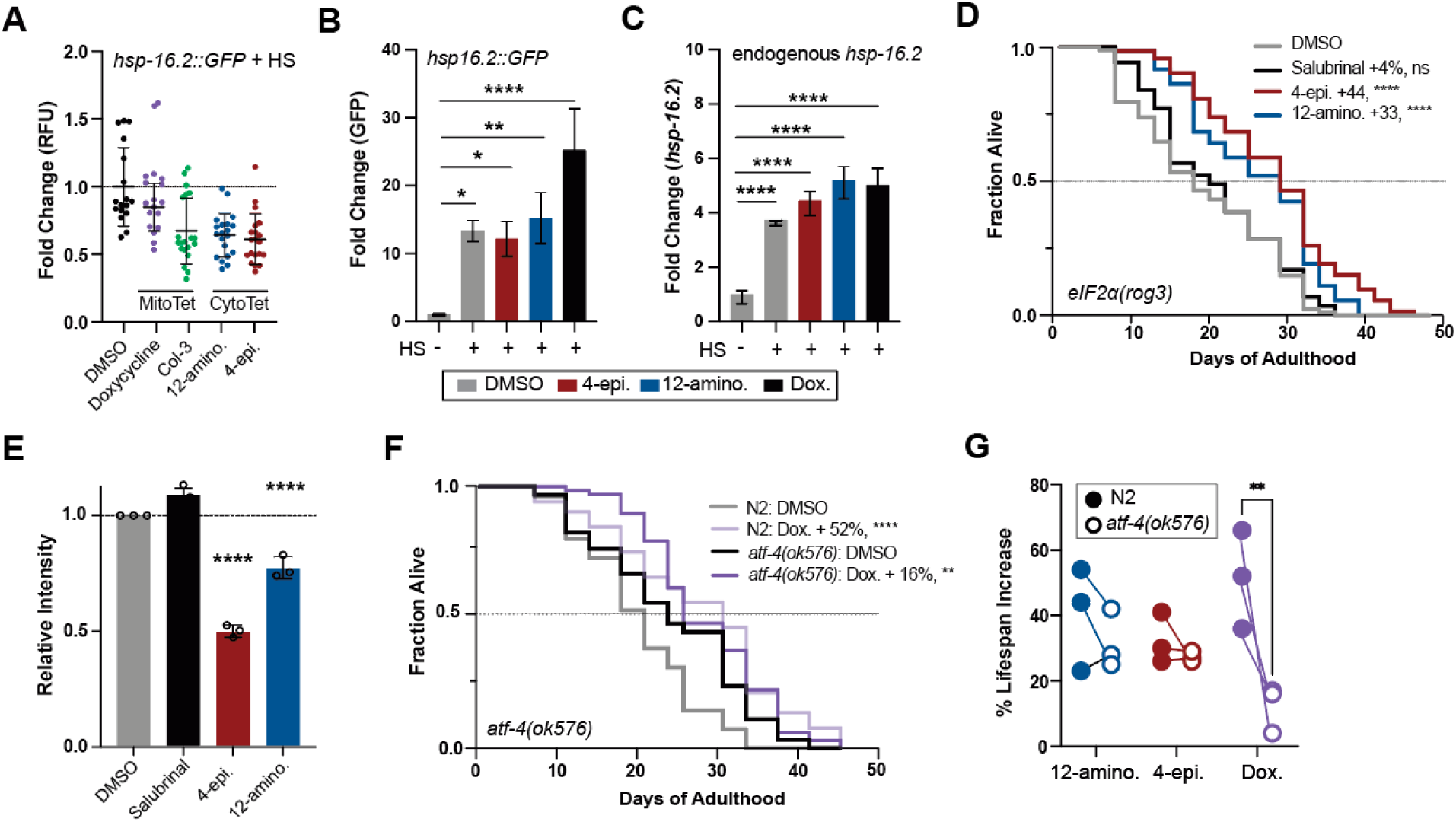
Tetracyclines elicit stress response-dependent and independent longevity mechanisms. **(A)** Scatter plot shows the fold induction relative to DMSO of the *hsp16.2::GFP* heat shock response reporter after tetracycline treatment, followed by a 1.5 hr, 35°C heat shock (HS)**. (B)** qRT-PCR quantification of relative GFP mRNA expression from untreated or tetracycline-treated animals with and without a 1.5 hr, 35°C HS. Tetracycline treatment suppresses the HS-induced GFP fluorescence of the *hsp-16.2p::GFP* reporter at the protein **(A)** but not at the mRNA level **(B)**. **(C)** qRT-PCR quantification of endogenous *hsp16.2* mRNA expression following HS with or without pre-incubation of tetracycline treatment of wild-type (N2) animals. **(D)** Survival plot of ISR-deficient *eIF2⍺(rog3)* mutants, which lack the eIF2*⍺* phosphorylation site. Only tetracyclines, but not salubrinal treatment, extend lifespan. **(E)** Quantification of 3 biological replicates from the SUNSET experiment in Figure 5F shows that 4-epiminocycline and 12-aminominocycline do not depend on *eIF2⍺* phosphorylation for translation inhibition, while salubrinal does. **(F)** Survival plot of N2 and the partially ISR-deficient *atf-4(ok576)* mutant treated with DMSO or doxycycline. ATF-4 is partially required for lifespan extension by doxycycline. The survival curve is one of the three trials quantified in Extended Figure 5G. **(G)** Comparison of the mean lifespan extension of N2 and *atf-4(ok576)* animals treated with the indicated tetracycline. Doxycycline specifically loses efficacy in *atf-4(ok576)* mutants across 3 biological replicates. Statistics for **B**, **C**, **G**: Significance was determined by one-way ANOVA with Dunnett’s multiple comparisons, where ** = p < 0.01, *** = p < 0.001, **** = p < 0.0001. All error bars indicate mean ± SD from three independent trials. Significance for all survival data (**D**, **E**) was determined by the log-rank test.

**Extended Figure 5:**
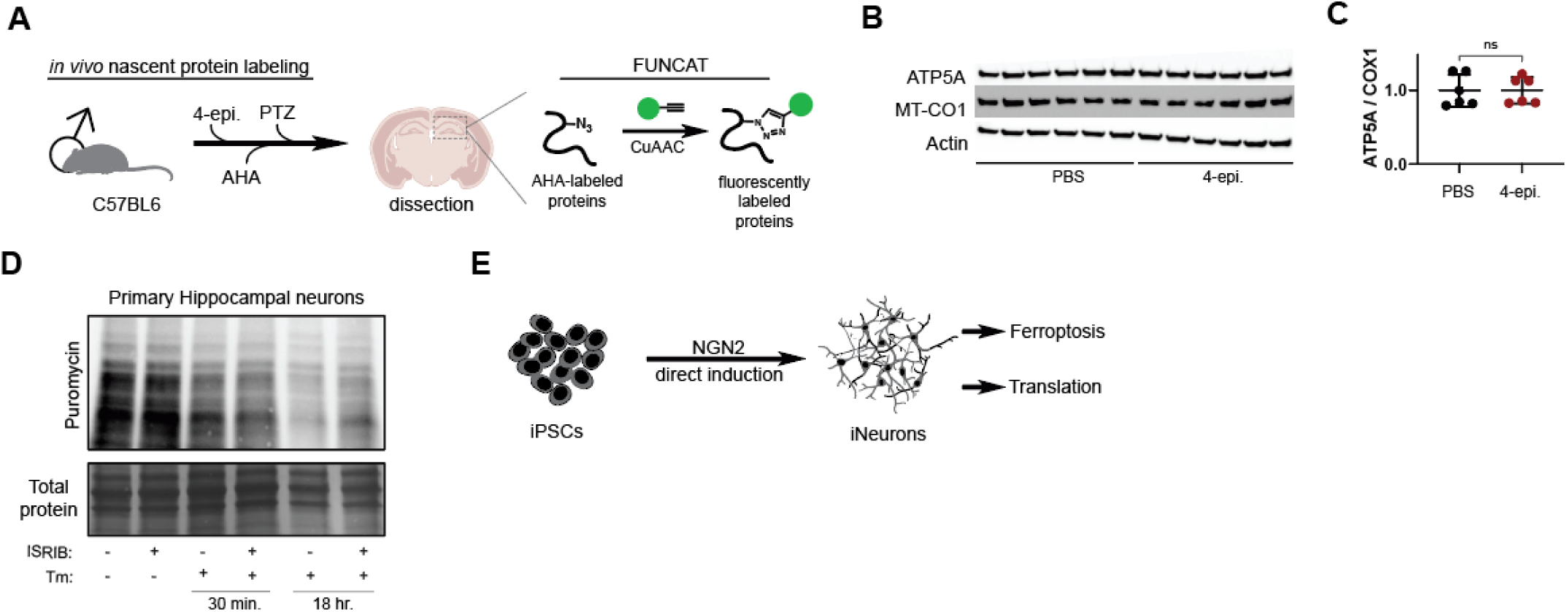
4-epiminocycline targets translation in mammalian neurons. **(A)** Experimental scheme to measure *de novo* translation by FUNCAT in the brains of male C57B6 mice. Following 4-epiminocycline i.p. injections (100 mg/kg), the animals were injected with AHA (50 mg/kg) and then with pentylenetetrazole (50 mg/kg) 30 minutes later. The hippocampus was micro-dissected and lysed to fluorescently label newly synthesized proteins that incorporated AHA by click chemistry. **(B)** 4-epiminocycline did not alter the ratio of nuclear–encoded (ATP-5A) to mitochondrial-encoded (MT-CO1) electron transport chain proteins. **(C)** Quantification of Extended Figure 5B. Actin was used as a loading control for normalization between samples before comparison of nuclear/mitochondrial–encoded proteins. ns = p > 0.05. **(D)** Validation of ISRIB activity in Figure 5E. Primary neurons were co-treated with 1 µM tunicamycin (Tm) to confirm the ability of ISRIB (300 nM) to rescue translation inhibited by tunicamycin. The 18 hr timepoint shows significant inhibition of translation by Tm, which is rescued with ISRIB. **(E)** Schematic outlining the generation of iNeurons from APOE3 (HE463#7) iPSCs and experimental procedure.

## Experimental Models

### Mouse studies

All procedures are approved by TSRI IACUC committee. C57BL/6 female mice (Stock #000664) were purchased from Jackson Labs at 9 weeks of age. Mice were group-housed and maintained on a 12 hrs light/dark cycle with ad libitum access to food and water.

### C. elegans strains

The Bristol strain (N2) was used as the wild-type strain. In addition, the following worm strains used in this study were obtained from the Caenorhabditis Genetics Center (CGC; Minneapolis, MN, USA): SJ4100 [zcIs13 [*hsp- 6p::GFP*]], CL2070 [dvIs70 [*hsp-16.2p::GFP + rol-6(su1006)*]], PS3551 [*hsf-1(sy441*)], RB790 [*atf-4(ok576)*], ZD1866 [*eIF2a(qd338)]*, ANR165 [*eIF2a(rog3)*]. Strains were backcrossed at least three times prior to experimental analysis.

### Cell Culture

HEK293 cells (ATCC CRL-1573) were directly obtained from ATCC. HT22 neuronal cells were obtained as a gift from Dr. Pamela Maher. Cells were grown in Dulbecco’s Modified Eagle Medium (DMEM, ThermoFisher, Cat#11995065) supplemented with 10% fetal bovine serum (FBS, ThermoFisher, Cat#16000044) and 1% penicillin-streptomycin (P/S, ThermoFisher, Cat#15140-122). Cells were subcultured every two or three days at 37 °C with 5% CO_2_ in a humidified incubator. Mycoplasma testing was performed every 6 months through TSRI.

Primary neurons were harvested as previously described in Chassefeyre et al^1^. Briefly, the hippocampi or cortices of male and female C57BL/6 P0-P1 mouse pups were dissected in Hanks’ balanced salt solution (HBSS) (ThermoFisher, Cat#24020117) supplemented with 0.08% d-glucose (Sigma-Aldrich, Cat#G6152), 0.17% HEPES (Sigma-Aldrich, Cat#H7006), and 1% P/S; filter-sterilized; and adjusted to pH 7.3. Dissected brain tissues were washed twice with cold HBSS and individually incubated at 37°C for 15 to 20 minutes in a sterile solution of 45 U of papain (Worthington, Cat#LS003119), 0.01% deoxyribonuclease (DNase) (Sigma-Aldrich, Cat#D4527), 1 mg of DL-cysteine (Sigma-Aldrich, Cat#C9768), 1 mg of bovine serum albumin (BSA) (Sigma-Aldrich, Cat#A7906), and 25 mg of D-glucose in phosphate-buffered saline (PBS) (ThermoFisher, Cat#10010049). Tissues were washed twice with DMEM supplemented with 10% FBS (preheated to 37°C) and disrupted by 10 to 12 cycles of aspiration through a micropipette tip. Dissociated neurons were then resuspended in warm DMEM supplemented with 10% FBS, counted and plated in either 96-well plates (Millipore Sigma, Cat#CLS3603) or 10 cm dishes (Genesee Scientific, Cat#25-202) pretreated with poly-L-lysine (50 μg/mL) (Sigma-Aldrich, Cat#P5899) in borate buffer [1.24 g of boric acid (Thermo Fisher Scientific, Cat#BP168-1) and 1.90 g of borax (Sigma-Aldrich, Cat#B9876) in 500 mL of cell culture–grade water, adjusted to pH 8.5, and filter-sterilized]. Plating densities were 2 x 10^4^ cells per well or 7.75 x 10^6^ cells per dish. After 3 hrs, medium was replaced with Neurobasal-A media (ThermoFisher, Cat#10888022), supplemented with 2% antioxidant free (ThermoFisher, Cat#10889038) or complete (ThermoFisher, Cat#17504044) B27 and 0.25% GlutaMAX (ThermoFisher, Cat#35050061). Cultures were maintained in culture media at 37 °C with 5% CO_2_. Four days after plating (DIV4), 5-Fluoro-2’-deoxyuridine (FUDR, Millipore Sigma, Cat#F0503) was added to a final concentration of 5 μM in culture media.

Induced neurons (iNeurons) were previously generated in Verduzco Espinoza et al^2^. In short, iNeurons were generated from APOE3/E3 (HE463#6) iPSCs, which is available on WiCell in a collection called Topol Lab’s Next Gen Cell Lines with the accession number of SCRP2307i. iPSCs were thawed in mTeSR Plus with 10 μM Y-27632 (Stem Cell Technologies, Cat#72304) and seeded on 6-well dishes coated with Matrigel. For maintenance, iPSCs were fed every 1-3 days, depending on confluence, and passaged 1-2 times per week as small clusters using 1 μM EDTA (Invitrogen, Cat#15575-020). 1X P/S was added to all culture media. Neurons were generated from iPSCs via direct induction with NGN2. iPSCs were dissociated with accutase (Stem Cell Technologies, Cat#7920) and seeded on Matrigel-coated 6-well plates at 1.5 x 10^5^ cells per well in mTeSR Plus with 10 μM Y-27632. The next day (day 0), each well was transduced with 125 μL of tetO-NGN2 and RTTA lentivirus for 2 hrs in mTeSR Plus with 10 μM Y-27632 at 37 °C. After 2 hrs, the viral media was aspirated and replaced with mTeSR Plus with 2 μg/mL doxycycline (Stem Cell Technologies, Cat#72742) to start NGN2 expression. On day 1, cells were fed with half mTeSR plus half Neuronal media [Neurobasal A, 1X B27, NEAA, Glutamax (ThermoFisher, Cat#10888022, 17504044, 350500661, 11140050] with doxycycline. NGN2-expressing cells were selected on day 2 by media exchange with doxycycline and puromycin (Gibco, Cat#A11138-03). On day 3, cells were replated at 6.25 x 10^4^ cells per cm^2^ on plates coated with PDL and Matrigel. 40 nM BRDU was added to select against the remaining dividing iPSCs. Cells underwent a full media change on days 7 and 10 to remove doxycycline and add neuronal differentiation factors (BDNF, GDNF, NT3 at 10 ng/mL, and laminin at 1 μg/mL). Experiments in this manuscript begin on day 14 for FUNCAT and day 24 for ferroptosis cell survival. For ferroptosis cell survival experiment, cells underwent half-media change on days 14 and 17.

### Compounds

Tetracyclines: Minocycline (MP Biomedicals, Cat#155718), 4-epiminocycline (Toronto Research Chemicals, Cat#TRC-E588540), Doxycycline (Sigma, Cat#D9891), Col-3 (Incyclinide, MedChemExpress, Cat#HY-13648), R464449 (Sigma-Aldrich, Cat#R464449), 12-aminominocycline (Toronto Research Chemicals, Cat#TRC-A618285), Tigecycline (LKT Laboratories, Cat#T3324). Other translation inhibitors: Cycloheximide (CHX, EMD Millipore, Cat#239763), Tunicamycin (Tm, Cayman Chemical, Cat#11445), Thapsagargin (Thaps., MedChemExpress, Cat#HY-13433). Following the initial hit, 4-epiminocycline and 12-aminominocycline were synthesized from minocycline to confirm purity and compound identity.

4-epiminocycline: **^1^H NMR** (400 MHz, DMSO-*d*_6_): *δ* 15.06 (s, 1H), 11.48 (s, 1H), 9.48-9.29 (m, 2H), 7.49-7.40 (m, 1H), 7.38 (s, 1H), 6.85-6.80 (m, 2H), 4.71 (s, 1H), 3.05-2.65 (m, 8H), 2.65-2.750 (m, 2H), 2.20-2.00 (m, 1H), 1.55-1.41 (m, 1H). **LCMS:** calc. for C_23_H_27_N_3_O_7_: 457.18, found: [M+H]^+^ 458.1. **HPLC:** 99.2% (254 nm). **SFC:** 97.7%.

12-aminominocycline: **^1^H NMR** (400 MHz, DMSO-*d*_6_): δ 13.17 - 13.03 (m, 1H), 10.72 - 10.13 (m, 1H), 9.69 - 9.46 (m, 1H), 9.10 - 8.44 (m, 1H), 7.16 (d, J = 8.8 Hz, 1H), 6.76 (d, J = 8.8 Hz, 1H), 6.46 - 6.21 (m, 1H), 5.95 (br s, 1H), 3.23 (dd, J = 4.0, 15.2 Hz, 1H), 2.98 (d, J = 1.6 Hz, 1H), 2.93 - 2.74 (m, 1H), 2.65 (br t, J = 2.4 Hz, 1H), 2.62 (t, J = 2.4 Hz, 1H), 2.57 (s, 6H), 2.47 (s, 4H), 2.39 (br s, 2H), 2.09 (br t, J = 14.4 Hz, 1H), 1.93 (br dd, J = 5.2, 7.6 Hz, 1H), 1.46 - 1.35 (m, 1H). **LCMS**: calc. for C_22_H_28_N_4_O_6_: 456.20, found: [M+H]^+^ 457.2. **HPLC:** 96.4% (254 nm).

## Method Details

### Lifespan assay

Age-synchronized *C. elegans* were prepared in liquid medium [S-complete medium with 50 mg/mL carbenicillin and 0.1 mg/mL fungizone in flat-bottom, optically clear 96-well plates (Corning, Cat#351172) containing 150 μL total volume per well, as previously described.^3^ Plates contained ∼10 animals per well in 6 mg/mL OP50. All experiments were performed with X-ray-irradiated OP50. Age-synchronized animals were seeded as L1 larvae and grown at 20 °C. Plates were covered with sealers to prevent evaporation. To prevent self-fertilization, FUDR (0.12 mM final) was added 42 – 45 hr. after seeding. Drugs were added on the days indicated and survival was scored manually by visualizing worm movement using an inverted microscope 3x/ week. When used, DMSO was kept to a final concentration of 0.33% v/v. Statistical analysis was performed using the Mantel–Haenszel version of the log-rank test as outlined in Petrascheck and Miller^4^.

### O-propargyl puromycin translation

HEK293 (6 x 10^5^) or HT22 (5 x 10^3^) cells were seeded into black 96-well plates (Costar, Cat#3603) with complete media (100 μL/well) and incubated overnight. For drug treatment, drugs were prepared at 20 mM stock in 100% DMSO. This was further diluted with 12.5% DMSO/PBS to make 12X working solutions at the indicated concentrations. 4 μL were added for a final concentration of 0.5% DMSO/PBS in each well. 0.5% DMSO/PBS served as a control. Drugs and control were added in triplicate and incubated for 2 hrs. Protein synthesis rates were determined using EZClick Global Protein Synthesis Assay Kit (BioVision, Cat#76305-296) according to the manufacturer’s protocol. In short, nascent polypeptides were pulse-labeled by addition of 1 μL 100X “protein label” and incubated for 30 minutes at 37 °C. We included a 1 μL addition of DMSO as a “no-protein label” control to subtract background fluorescence. Then culture media was removed and cells were washed once with 100 μL PBS. Cells were fixed by addition of 100 μL “fixative solution” and incubated for 15 minutes on ice in the dark. This was aspirated off and 100 μL “permeabilization buffer” was added and incubated for 10 minutes at room temperature. Buffer was removed and 20 μL fresh “permeabilization buffer” was added. The 1X EZclick reaction cocktail was prepared according to the protocol: 3 μL PBS, 1 μL copper reagent, 1 μL fluorescent azide, 5 μL reducing agent, scaled for number of reactions needed. 100 μL 1X EZclick reaction cocktail was added to each well and incubated. After 30 minutes, the reaction mixture was aspirated and cell were washed three times with 100 μL “wash buffer.” Wells were then incubated with 1X DAPI stain for 15 minutes and washed three times with “wash buffer”. As a positive control, 0.5 μM cycloheximide (CHX) was added 30 minutes prior to addition of “protein label.” When thapsagargin was used, it was added at the same time as the tetracyclines at 250 nM.

Each well was imaged using the ImageExpress HT.ai confocal high-content imaging system (IXM, Molecular Devices). A 20X Apo LWD 9na 0.95 water immersion objective was used with a confocal 50 μm slit setting. Laser power was set to 2% and each image frame was exposed for 100ms for GFP channel and 10ms for DAPI channel. Each well was imaged in 4 center locations. Images were analyzed using a custom Cell Profiler pipeline that autodetects an ROI based on DAPI staining of nuclei, then expands ROI by 20 pixels, and then measures GFP signal. Measuring GFP/DAPI signal gives an approximation of total protein synthesis relative to total cells seeded in each well.

### Oxytosis Ferroptosis survival assay—HT22 cells

4 x 10^3^ cells were seeded into sterile black 96-well plates with complete media and incubated overnight (100 μL/well). For drug treatment, drugs were prepared at 10 mM stock in 100% DMSO. This was further diluted with 5% DMSO/PBS to make 25X working solutions. 4 μL were added for a final concentration of 0.5% DMSO in each well. 0.2% DMSO served as a control. Drugs and control were added 2 hrs prior to addition of oxytosis/ferroptosis inducers. 5 mM glutamate (ThermoFisher, Cat#156212500) was added to each well and cell viability was measured using CellTiter-Glo (Promega, Cat#G9243). Luminescence was measured using BioTek Cytation 5 (Agilent). Results are shown as a percentage of untreated control cells. In experiments where ISRIB was used to block the ISR, 300 nM ISRIB was added 2 hrs prior to the addition of drugs/DMSO controls. When thapsagargin was used, it was added at the same time as the tetracyclines at 250 nM.

### Oxytosis Ferroptosis survival assay—Primary and iNeurons

For primary neurons: Hippocampal primary neurons were prepared in sterile black 96-well plates coated with borate buffer as described above. At DIV7, drugs were added at increasing concentrations to each well and incubated for 4 hrs. Then, 300 nM RSL3 (Selleck Chem, Cat#S8155) was added to each well and incubated for 24 hrs. Viability was determined using CellTiter-Glo as above. When ISRIB was used, 300 nM was added to each well and incubated for 4 hrs prior to drug/control treatment.

For iNeurons: On day 3, 24 hrs after puromycin selection, cells from a 6-well plate were replated equally into 96-well plates, at 2 x 10^4^, leaving the edge wells empty. 40 nM BRDU was added to select against the remaining dividing iPSCs. The media was changed on days 7 and 10 with neuronal differentiation factors (BDNF, GDNF, NT3, Laminin). Cells were maintained with half media changes twice a week and were then allowed to grow until day 24. Then 0, 0.14, 0.41, 1.2, 3.7, 11, 33, or 100 μM 4-epiminocycline was added to 6 wells of the 96-well plate. After 2 hrs, 2 μM RSL3 was added to each well to induce ferroptosis. 24 hrs later, viability was determined using CellTiter Glo as above.

### Surface Sensing of Translation (SUNSET)

For *C. elegans*: Day 1 adult N2 worms were bleached, and eggs were allowed to hatch in S-complete by shaking overnight. On the next day, 10,000 L1 worms were seeded in a 10 cm plate containing a total volume of 20 mL S-complete with 6 mg/mL OP50 bacteria, 50 μg/mL carbenicillin, and 0.1 μg/mL amphotericin B. 4 mL FUDR (0.6 mM stock in S-complete) were added to worms at L4 stage in each plate. 2 hrs later, 100 μM of each compound was added to worms. After 12 hrs, worms were transferred into a 15 mL corning tube containing a total volume of 4 mL S-complete with 500 μL 6 mg/mL OP50 bacteria, 0.5 mg/mL puromycin (ThermoFisher, Cat#A11138-03), and 100 μM tetracyclines. After rotating the corning tubes for 4 hrs, worms were collected into 2 mL cryotubes by washing them with M9 once and with cold PBS three times. Worms were flash-frozen in liquid nitrogen and 150 µL of cold lysis buffer [20 mM Tris base, 100 mM NaCl, 1 mM MgCl_2_, pH = 7.4, with protease inhibitors (Roche, Cat#11836153001)] was added and samples subsequently broken with a beak mill homogenizer (Fisher). Protein concentrations were determined by the Bradford protein assay (Bio-Rad, Cat#5000006). 50 μg protein from each sample was loaded for western blot analysis using antibodies against puromycin (Millipore, Cat#MABE343) and GAPDH (Proteintech, Cat#10494-1-AP). Primary antibodies were diluted 1:5,000 in 5% non-fat milk in TBST, and secondary antibodies were diluted 1:10,000. To determine the relative intensities of each blot, the integrated intensity was measured for each full anti-puromycin lane using ImageJ. A similar sized band with no signal was used to calculate and subtract background, and then each intensity was normalized to corresponding GAPDH loading control. Finally, each condition was normalized to DMSO control for quantification and statistics.

For primary neurons: Cortical primary neurons were prepared in 10 cm dishes as described above. On DIV7, dishes were treated with 300 nM ISRIB or DMSO. Immediately after, tunicamycin was added to a final concentration of 3 ug/mL. Either 30 minutes or 18 hrs after tunicamycin addition, puromycin was added to a final concentration of 2ug/mL. 2 hrs after puromycin addition, total protein was extracted with RIPA buffer [150 mM NaCl (Sigma-Aldrich, Cat#S7653), 5 mM EDTA (Applichem, Cat#A1104), 50 mM Tris (ThermoFisher, Cat#BP152-5), 1% NP-40 (Millipore Sigma, Cat#492016), 0.5% sodium deoxycholate (ThermoFisher, Cat#BP349-100), and 0.1% SDS] supplemented with protease and phosphatase inhibitor cocktail (ThermoFisher, Cat#78442). Protein concentrations were determined by the Bradford protein assay. 15 μg from each sample was loaded for western blot analysis using antibodies against puromycin diluted 1:1000 in 5% non-fat milk in TBST. Relative intensities were determined as described above. Total protein was visualized with Coomassie stain solution (0.1% Coomassie Brilliant Blue R stain (ICN Biomedicals, Cat#02190343-CF), 7% acetic acid, 50% methanol).

### Antibiotic activity assay

One colony of OP50 was inoculated overnight in a 37 °C shaker in 5 mL LB media containing 50 μg/mL ampicillin. 14-16 hrs later, the pre-inoculum was diluted 1:200 in 5 mL LB containing 50 μg/mL ampicillin. In a flat bottom, clear 96-well plate, 100 μL of the diluted inoculum was added to each well being tested, one column (8 wells) per condition. DMSO or the indicated drugs were added and a 0 time point was recorded for baseline OD_600_ values using a plate reader (Tecan Safire II). The plate was returned to the 37°C shaker and OD_600_ was re-measured every hr. OD_600_ values for each time point were averaged across each well per condition.

### Zn^2+^ chelation assay

Chelation assay was preformed using the Abchem Zinc Assay Kit (Abcham, Cat#ab102507). First, the included zinc standard was added to each well according to the manufacturers protocol. In a clear 96-well plate, samples were preincubated with 0, 100, 300, 780, 1200, 1800 μM of each tetracycline in duplicate. Samples were allowed to incubate for 30 minutes before determination of free Zn^2+.^. Absorbance was measured at OD_560nm_ on the Tecan Safire II. EDTA was included as a positive control and data was normalized with 0% remaining free Zn^2+^ defined by assay buffer with no zinc standard, and 100% free zinc being the DMSO control.

### MMP9 Inhibition

MMP9 inhibition was determined using a quenched fluorogenic peptide included in the MMP9 Inhibitor Screening Assay Kit following the manufacturer’s protocol (Abcham, Cat#ab139449). N-hydroxy-2-phenylethanamide (NNGH, 1 μM) was included as a positive control. In the screening assay, 100 μM of each tetracycline in triplicate was added to a well of a 96-well plate containing MMP enzyme and assay buffer and allowed to incubate for 10 minutes. MMP9 substrate was then added to each well using a multi-channel pipet, the plate was incubated for 60 minutes at 37 °C, and fluorescence was measured using the BioTek Cytation 5. An identical procedure was followed for the generation of dose response curves for 12-aminominocycline, 4-epiminocycline, minocycline, and tigecycline except concentrations were: 0, 1, 10, 30, 100, 300 μM. 4 technical replicates were used for each concentration and assay was repeated twice.

### UPR^MT^ stress imaging

20,000 SJ4100 [zcIs13 [*hsp- 6p::GFP*]], age synchronized animals were prepared as in the SUNSET method. At the developmental stage of L2, 100 μM of each tetracycline or DMSO was added and allowed to incubate for 12 hrs. After the treatment window, animals were washed 3x with S-complete to remove any bacteria and suspended in 10 mL S-complete. GFP intensity was quantified using the COPAS Biosorter (Union Biometric). Integral values for GFP channel were used to determine fluorescent values. The animals were also sorted into 96-well plates and imaged using an ImageXpress Micro XL High-Content screening system (Molecular Devices) with a 2x objective.

### HSR stress imaging

20,000 CL2070 [*dvIs70 [hsp-16.2p::GFP+rol-6(su1006)]]* animals were prepared as in the SUNSET method. 100 μM of each tetracycline or DMSO was added to liquid culture at the late L4 stage. On day 1 of adulthood, after 12 hrs treatment, the liquid culture was transferred to a 50 mL falcon tube and worms were settled by gravity. The supernatant was aspirated off leaving the concentrated worm pellet. The pellet was washed twice with S-complete, pelleted by gravity again, and then transferred to 10 cm NGM plate. Once the liquid was completely dry on the NGM plate, the plates were transferred to a 36 °C incubator, upside down, and incubated for 1.5 hrs. Following heat shock, NGM plates were transferred to 20 °C incubator and allowed to recover for 8 hrs. Worm GFP intensity was determined as in the *hsp-6::GFP* reporter using the COPAS Biosorter, but instead of being sorted into 96-well plates, 5,000 animals were bulk-sorted into 2 mL cryotubes and flash frozen for RT-qPCR analysis.

### Quantitative real-time PCR (qRT-PCR) and data analysis

All qRT-PCR experiments were conducted according to the MIQE guidelines, except that samples were not tested in a bio-analyzer, but photometrically quantified using a Nanodrop. Worm pellets were obtained from the HSR stress imaging experiment above. To extract RNA, frozen animals were suspended in ice-cold Trizol (Qiagen, Cat#79306), then zirconium beads were added and animals were broken open with a beak mill homogenizer (Fisherbrand). Following chloroform extraction, RNA was precipitated using isopropanol and washed once with 75% ethanol followed by DNAse (Sigma-Aldrich, Cat#AMPD1-1KT) treatment. Reverse transcription was carried out using iScript RT-Supermix (Bio-Rad, Cat#170–8841) at 42°C for 30 minutes. Quantitative PCR reactions were set up in 384-well plates (Bio-Rad, Cat#HSP3901), which included 2.5 µl Bio-Rad SsoAdvanced SYBR Green Supermix (Cat#172–5264), 1 µl cDNA template (2.5 ng/µl, to final of 0.5 ng/µl in 5 µl PCR reaction), 1 µl water, and 0.5 µl of forward and reverse primers (150 nM final concentration). Quantitative PCR was carried out using a Bio-Rad CFX384 Real-Time thermocycler (95°C, 3 minutes; 40 cycles of 95°C 10 s, 60°C 30 s; Melting curve: 95°C 5 s, 60–95°C at 0.5°C increment, 10 s). Gene expression was normalized to three reference genes for *C. elegans* samples, *act-1*, *xpg-1* and *rpl-6*.

### Fluorescent Non-canonical Amino Acid Tagging (FUNCAT)

For iNeurons: On day 3, 24 hrs after puromycin selection, cells from a 6-well plate were replated equally into 6-well palates, at 6 x 10^5^. on day 14, 0, 10, 30, or 100 μM 4-epiminocycline was added to each well of a 6 well plate (3 mL media). 250 nM CHX was added as a positive control in one of the wells. Drugs were prepared from 30 mM stock solutions in 100% DMSO. Working solutions were prepared at 300X in 15% DMSO and 20 μL was added to each well for a final concentration of 0.1% DMSO and allowed to incubate for 2 hrs. 200 mM azidohomoalanine (AHA, Vector Laboratories, Cat#CCT-1066-1000) stock solution was prepared with Ultrapure H_2_O and brought to pH = 7 with 10N NaOH. 200 mM AHA was diluted in complete culture media pre-warmed to 37 °C for a final concentration of 8 mM. Media was replaced with AHA and pulse labeled for 2 hrs after which labeling was stopped by removal of AHA media.

iNeurons were washed once 1X DBPS and then lysed in dish with the addition of 100 μL 10% RIPA/DPBS buffer + 1X protease inhibitors (Pierce Protease Inhibitor Mini Tablets, EDTA-free, ThermoFisher, Cat#A32955) on ice. Cells were scrapped into an Eppendorf tube and centrifuged at 5000 x g for 2 minutes to remove debris. Lysate was collected and protein concentration was determined by Pierce BCA assay kit (ThermoFisher, Cat#23235) following manufacturer’s instructions.

Cell lysates were adjusted to 1.0 mg/mL using cold DPBS. To each sample (50 μL) in a PCR Eppendorf tube, 6 μL of freshly prepared click reaction mixture was added. The cocktail was prepared by addition of 3 μL THPTA (1.7 mM in H_2_O), 1 μL CuSO_4_ (50 mM in H_2_O), 1 μL TAMARA-azide (1.25 mM in DMSO),and 1 μL ascorbate (50 mM in H_2_O, prepared last). After addition of the mixture, each reaction was vortex, briefly centrifuged, and incubated at 40 °C for 1 hr. The reaction was quenched by addition of addition of 17 μL of 4X SDS loading buffer (Bio-Rad, Cat#1610747), and heated at 95 °C for 5 minutes. 30 μg protein was loaded per lane of a polyacrylamide gel (Bio-Rad, Cat#4569034) and visualized by in-gel fluorescence on a ChemiDoc MP flatbed fluorescence scanner (Bio-Rad, Cat#12003154). After imaging, gels were stained with Coomassie blue to determined total protein. Gels were quantified in ImageJ by dividing the integrated intensity of each lane of the fluorescent image by the integrated intensity of the corresponding total protein lane after subtracting a similar area background lane, then normalizing to the DMSO control. Abbreviations: THPTA ((tris-hydroxypropyltriazolylmethyl)amine), Combi-blocks, Cat#QH-3278) CuSO_4,_ (Copper(II) sulfate, Sigma-Aldrich, Cat#C1297), Azide-fluor 545 (5-carboxytetramethylrhodamine-azide, Millipore Sigma, Cat#760757), and ascorbate ((+) Sodium L-ascorbate, Sigma-Aldrich, Cat#A7631).

For *in vivo* translation: We utilized the *in vivo* translation measurement method developed in Xie et al^5^ but adapted for gel-based FUNCAT. 10-week old male C57BL6 were treated with either 4-epiminocycline or control in drinking water (0.6 mg/mL) for 3 days (n = 6 per group). This dose was selected because it is well tolerated in mice and we determined a plasma level of approximately 7 μM for the analog minocycline, which is comparable to circulating levels in human patients of 2–11 μM^6,7^. On the 4th day, 100 mg/kg 4-epiminocycline or control (saline) was administered via intraperitoneal (i.p.) injection. 30 minutes later, AHA (dissolved in PBS) was i.p. injected at 50 mg/kg. Again, 30 minutes later 50 mg/kg pentylenetetrazol (PTZ, dissolved in PBS) was i.p. injected. 90 minutes later the animals were sacrificed, PBS-perfused, the hippocampus was harvested, and samples snap frozen in liquid nitrogen.

Samples were resuspended in 150 μL 1x DPBS and homogenized with a hand-held tip sonicator (Fisher Scientific, sonic dismembrator model 100). Protein concentration of the homogenate was determined by BCA protein assay kit. Protein lysate with AHA-labeled nascent proteins was normalized by total protein to 2.0 mg/mL and the click reaction with TAMRA-azide was performed as described above.

### Rat pharmacokinetic determination

Pharmacokinetic testing for 4-epiminocycline was conducted by Pharmaron. 4-epiminocycline was formulated in PBS and was i.p. injected at a low dose of 25 mg/kg and high dose of 50 mg/kg. A total of 3/3 male rats were assessed at 0.083, 0.25, 0.5, 1, 2, 4, and 8 hr timepoints for blood. At the 8 hr timepoint, the mice were sacrificed after blood collection, and brains were weighed then homogenized. Analytes from each collection were extracted and run on an LC-MS/MS (AB API 55000) to quantify compound concentration. Data were collected as plasma (ng/mL) and brain (ng/g) concentrations determined by an internal standard method.

